# Single Trial Decoding of Movement Intentions Using Functional Ultrasound Neuroimaging

**DOI:** 10.1101/2020.05.12.086132

**Authors:** Sumner L. Norman, David Maresca, Vasileios N. Christopoulos, Whitney S. Griggs, Charlie Demene, Mickael Tanter, Mikhail G. Shapiro, Richard A. Andersen

**Affiliations:** California Institute of Technology, Pasadena, CA, USA; Physics for Medicine Paris, Inserm, CNRS, ESPCI Paris, PSL Research University, Paris, France; INSERM Technology Research Accelerator in Biomedical Ultrasound, Paris, France; T&C Chen Brain-Machine Interface Center at Caltech

**Author notes:** These authors contributed equally to this work (Sumner L. Norman, David Maresca, Vasileios N. Christopoulos). These senior authors contributed equally (Mikhail G. Shapiro, Richard A. Andersen). University of California: Riverside, Riverside, California, USA. ***Corresponding author(s)***, Correspondence should be addressed to M.G.S. or R.A.A.

## Abstract

Brain-machine interfaces (BMI) are powerful devices for restoring function to people living with paralysis. Leveraging significant advances in neurorecording technology, computational power, and understanding of the underlying neural signals, BMI have enabled severely paralyzed patients to control external devices, such as computers and robotic limbs. However, high-performance BMI currently require highly invasive recording techniques, and are thus only available to niche populations. Here, we show that a minimally invasive neuroimaging approach based on functional ultrasound (fUS) imaging can be used to detect and decode movement intention signals usable for BMI. We trained non-human primates to perform memory-guided movements while using epidural fUS imaging to record changes in cerebral blood volume from the posterior parietal cortex – a brain area important for spatial perception, multisensory integration, and movement planning. Using hemodynamic signals acquired during movement planning, we classified left-cued vs. right-cued movements, establishing the feasibility of ultrasonic BMI. These results demonstrate the ability of fUS-based neural interfaces to take advantage of the excellent spatiotemporal resolution, sensitivity, and field of view of ultrasound without breaching the dura or physically penetrating brain tissue.

## Main

Brain-machine interfaces (BMI) are a technology that provides a direct link between mind and machine by interpreting neural activity into user intention. They allow the user to control neuroprosthetic devices, including computer cursors and prosthetic limbs^1,2^. Recent studies have enabled people with paralysis to control prosthetic devices with bandwidth approaching that of unimpaired people^3^. BMI thus promises the ability to assist or restore movement to people living with severe paralysis. The most advanced neural interfaces are based on intracortical electrophysiology, which provides direct access to the electrical signals of neurons with excellent temporal resolution^1,4^. However, the electrodes used in these techniques must be implanted via high-risk open-brain surgery. This process causes acute and chronic tissue damage^5^ and implants suffer material degradation. These factors limit longevity and performance^6–8^. In addition, invasive electrodes are difficult to scale and limited in sampling density and brain coverage. Non-invasive electroencephalography (EEG) has been developed as a method for BMI since 1973^9^ and achieved considerable success in the research setting^10–16^. However, it is intrinsically limited by its topographical representation of the activity of large brain volumes and the dispersion of signal by volume conduction through various tissues and bone.

Here, we introduce the possibility of BMI based on functional ultrasound (fUS) imaging – a recently developed minimally invasive neuroimaging technique. fUS imaging is a hemodynamic technique that visualizes regional changes in blood volume using ultrafast Doppler angiography^17–19^. It provides excellent spatiotemporal resolution (< 100 μm and 100 ms) and high sensitivity (~ 1mm/s velocity^20^) across a large field of view (several cm). Since its introduction in 2011^17^, fUS has been used to image neural activity during epileptic seizures^17^, olfactory stimuli^21^, and behavioral tasks in freely moving rodents^22,23^. It has also been applied to many non-rodent species including ferrets^24^, pigeons^25^, non-human primates (NHPs)^26^, and humans^27–29^. Unlike functional magnetic resonance imaging (fMRI), fUS can be applied in freely moving subjects using miniature, head-mountable transducers. In addition, the hemodynamic imaging performance of fUS is approximately 5 to 10-fold better in terms of spatiotemporal resolution and sensitivity compared to fMRI, providing access to qualitatively different information.

In this study, we leveraged the exceptional sensitivity of fUS imaging to detect movement intentions in NHPs. We trained two animals to perform memory-guided saccades to peripheral targets and one animal to perform memory-guided reaches. We recorded neurovascular activity over the posterior parietal cortex (PPC) through a minimally invasive cranial window throughout the task. PPC is an association region situated between visual and motor cortical areas and involved in high-level cognitive functions including spatial attention^30^, multisensory integration^31^, and sensorimotor transformations for movement planning^32^. PPC contains several sub-regions with anatomical specialization. These include the lateral intraparietal area (LIP) involved in the planning of eye movements^31,33–35^ and the parietal reach region (PRR) involved in planning of reach movements^31,35–38^. The functional characteristics of PPC have been thoroughly studied using electrophysiological and magnetic resonance techniques^32,37,39–42^, providing ample evidence for comparison to the fUS data. Importantly, LIP and PRR are adjacent areas and can be captured synchronously within a single fUS imaging frame (approximate slice volume of 12.8 x 13 x 0.4 mm). Finally, because PPC encodes high level aspects of movement planning, it presents a unique source of pre-movement and goal-related control signals for BMIs, an important factor when considering hemodynamic recording methods. By recording brain activity while animals perform movement tasks, we provide the first evidence that fUS can detect motor planning activity that precedes motor responses in NHPs. Importantly, we extend these results in the context of a recording method for BMIs, performing offline decoding of the intended movement direction of the animals on a single-trial basis. These results present for the first time a proof of concept that fUS is capable of imaging neurovascular dynamics in large animals with enough sensitivity to decode the goal of an intended movement. fUS is therefore a viable recording technique for brain-machine interfacing.

## Results

To look for goal-related neurovascular signals in the PPC, we acquired fUS images from NHPs using a miniaturized 15 MHz, 128-element linear array transducer inserted epidurally through a cranial window. The transducer provided a spatial resolution of 100 μm x 100 μm in-plane, slice thicknesses of ~400 μm, covering a plane with a width of 12.8 mm and a penetration depth of 16 mm. We positioned the probe surface-normal in a coronal orientation on the dura above the PPC (**Fig. 1, a-b**). We then selected planes of interest for each animal from the volumes available (**Fig. 1, c-f**). Specifically, we chose planes that exhibited behaviorally tuned hemodynamic activity and captured both lateral and medial banks of the intraparietal sulcus (IPS) within a single image, allowing us to assess the roles of multiple brain regions synchronously. The geometry of ultrasound imaging made it convenient for us to access cortical regions in sulci. Within the chosen planes, we captured ultrafast power Doppler images at a frame rate of 1 Hz.

**Figure 1:**
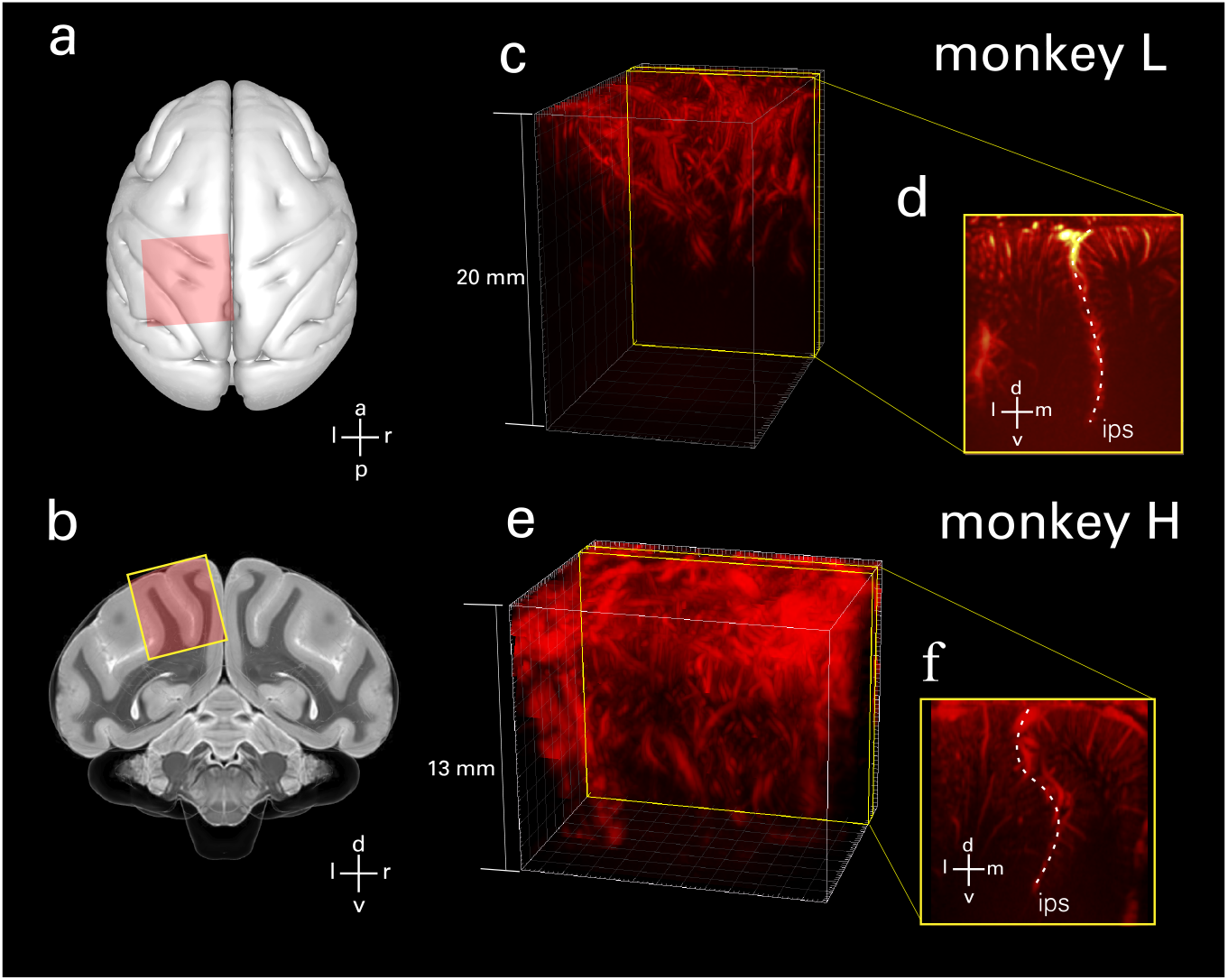
**a-b**, Anatomical illustrations of craniotomy field of view in an axial plane (**a**) and a coronal cross-section (**b**) overlaid on a NHP brain atlas^43^. The 24×24mm (inner dimension) chambers were placed surface normal to the brain on top of the craniotomized skull. 3D vascular maps for each of the two NHPs: monkey L (**c**) and monkey H (**e**). The field of view included the central and intraparietal sulci for both monkeys. A representative slice for monkey L (**d**) and monkey H (**f**) shows the intraparietal sulcus (dotted line, labeled) with orientation markers (l=lateral or left, r=right, m=medial, v=ventral, d=dorsal, a=anterior, p=posterior).

### Hemodynamic response during memory-guided saccades

To resolve goal-specific hemodynamic changes within single trials, we trained two NHPs to perform memory-delayed saccades. We used a similar task design to previous experiments investigating the roles of PPC regions using fMRI BOLD^39,44^ and pharmacological inactivation^45^. Specifically, the monkeys were required to memorize the location of a cue presented in either the left or right hemifield and execute the movement once the center fixation cue extinguished (**Fig. 2a**). The memory phase was chosen to be sufficiently long (5.1 sec +/- 1.4 sec) to capture transient hemodynamics. We collected fUS data while each animal (N=2) performed memory-delayed saccades. In total, we collected data for 30 runs over 18 days. Each day consisted of up to five runs, where each run included 30-minutes of randomly generated targets for one task.

**Figure 2:**
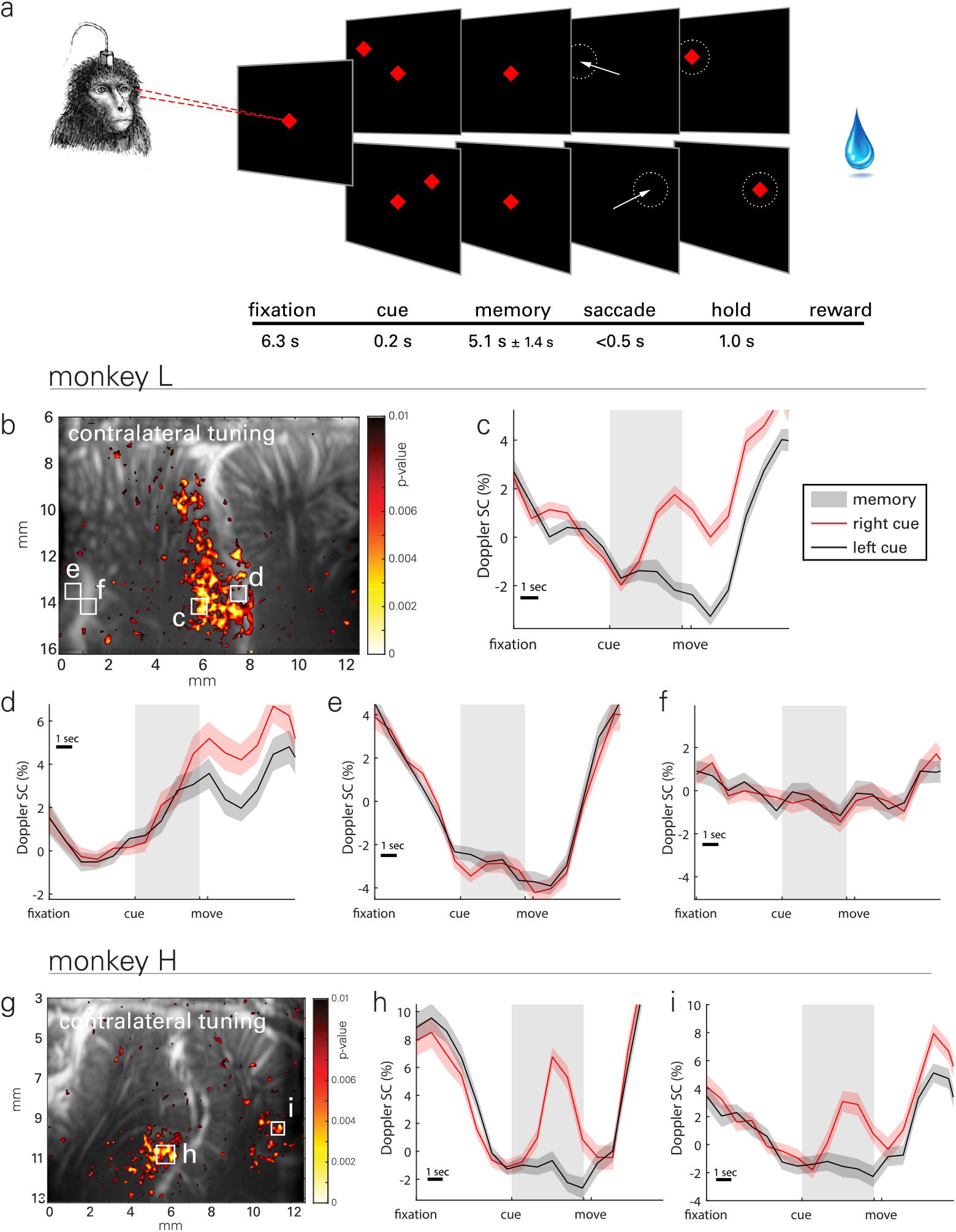
**a**, Memory-guided saccade task. A trial started with the animals fixating on a central cue (red diamond). Next, a target (red diamond) was presented either on the left or the right visual field. The target disappeared after 0.2 s and the animals had to remember its location while continuing to fixate on the center cue. When the cue was extinguished (go-signal), the animals performed a saccade to the remembered peripheral target location within 0.5 s and maintained eye fixation for another 1 s before receiving a reward. **b-f**, Representative activity map and event-related average (ERA) waveforms of CBV change during memory-guided saccades for monkey L. **g-i**, Activity map and ERA waveforms for monkey H. **b**, A statistical map shows localized areas with significant differences (t-test, p<0.01) in signal change (SC) during the memory delay phase for left vs. right cued saccades, i.e. vascular patches of contralaterally tuned activity. **c, d**, ERA waveforms in LIP display lateralized tuning specific to local populations. **e**, Small vasculature outside of LIP exhibits event related structure that is tuned to task structure but not target direction. **f**, Vessels that perfuse large areas of cortex do not exhibit event related signal. **g**, Map for monkey H. **h**, ERA waveforms show lateralized tuning in LIP. **i**, Target tuning also appears in medial parietal area (MP) for monkey H. Panels c-f share a common range (9% signal change) as do h-i (14%).

We use statistical parametric maps based on the Student’s t-test (with false discovery rate correction, see Materials and Methods) to visualize spatiotemporal patterns of lateralized activity in PPC (**Fig. 2b**). We observed event-related average (ERA) changes of CBV from localized regions as small as 100 x 100 μm (**Fig. 2, c-f**). We also present a representative map (**Fig. 2g**) and ERAs (**Fig. 2, h-i**) for the second animal. Spatial response fields of laterally tuned neurovascular activity were observed on the lateral bank of IPS, i.e. in LIP, for both animals, and ERA waveforms were similar between animals in this area. These response fields and waveforms are consistent with previous electrophysiological^46^ and fMRI BOLD^44^ results. Specifically, ERAs from this area show higher memory-phase responses to contralateral (right) compared to ipsilateral (left) cued trials (area under curve for memory phase, t-test p<0.001, **Fig. 2c**).

High sensitivity and resolution afforded by fUS allows us to distinguish the function between neighboring regions on the scale of hundreds of microns, e.g. within PPC sub-regions such as LIP. As an example, a second LIP region (**Fig. 2d**) also shows contralateral tuning, but the difference in activity occurs later in the memory period and persists into the movement period and inter-trial interval. One animal, monkey H, exhibited a similar direction-tuned response in the medial parietal area (MP) on the medial wall of the hemisphere. We did not record this area-effect in monkey L as we optimized recording location for activation in putative LIP rather than area MP; thus, area MP was outside the imaging plane. However, this tuning supports evidence of MP’s role in directional eye movement observed in a previous study^47^. In contrast, focal regions of microvasculature outside area LIP also showed strong event-related responses but were not typically tuned to target direction. In one example, a focal patch of the lateral aspect of the image (**Fig. 2e**) exhibited a large event-related response: a decrease in activity during the fixation period, consistent signal during the memory period and a sharp increase after movement. Notably, this response was consistent regardless of target direction.

### Single trial decoding - feasibility of a fUS-based BMI

The goal of a BMI is to predict the existence and parameters of an upcoming intended movement directly from the brain. To analyze the feasibility of fUS imaging for future BMIs, we set out to predict the direction of an upcoming eye movement on a single-trial basis. To demonstrate the robust nature of fUS signals, we present results from classifiers based on linear techniques of dimensionality reduction and class discrimination. We present a graphical representation of the decoding analysis in **Fig. 3a**. Briefly, we used classwise principal component analysis (CPCA) to reduce data dimensionality while maintaining enough components to retain >95% of the variance in the data. We then use ordinary least squares regression (OLSR) to regress the PCA-transformed data from the memory delay period to the movement direction, i.e. class label, and stored the resulting transformation matrix. For the testing set, we transformed the raw data from the memory delay period of each trial using the same PCA subspaces and OLSR transformation. Finally, we used linear discriminant analysis (LDA) to classify the resulting value for each trial as a presumed left or right movement plan. We used CPCA because it is ideally suited to discrimination problems with high dimension and small sample size, like ours^48,49^. All reported results were generated using a 10-fold cross validation. Successful trials varied from 59 to 387 trials in a given session; thus, we trained the model using between 53 and 348 trials on any given day. Saccade goal prediction accuracy within a session, i.e. decoded from the memory delay, ranged from 61.5% (binomial test vs. chance level, p=0.012) to 100% (p<0.001) on a given 30-minute run. The mean accuracy across all sessions and runs was 78.6% (p<0.001).

**Figure 3:**
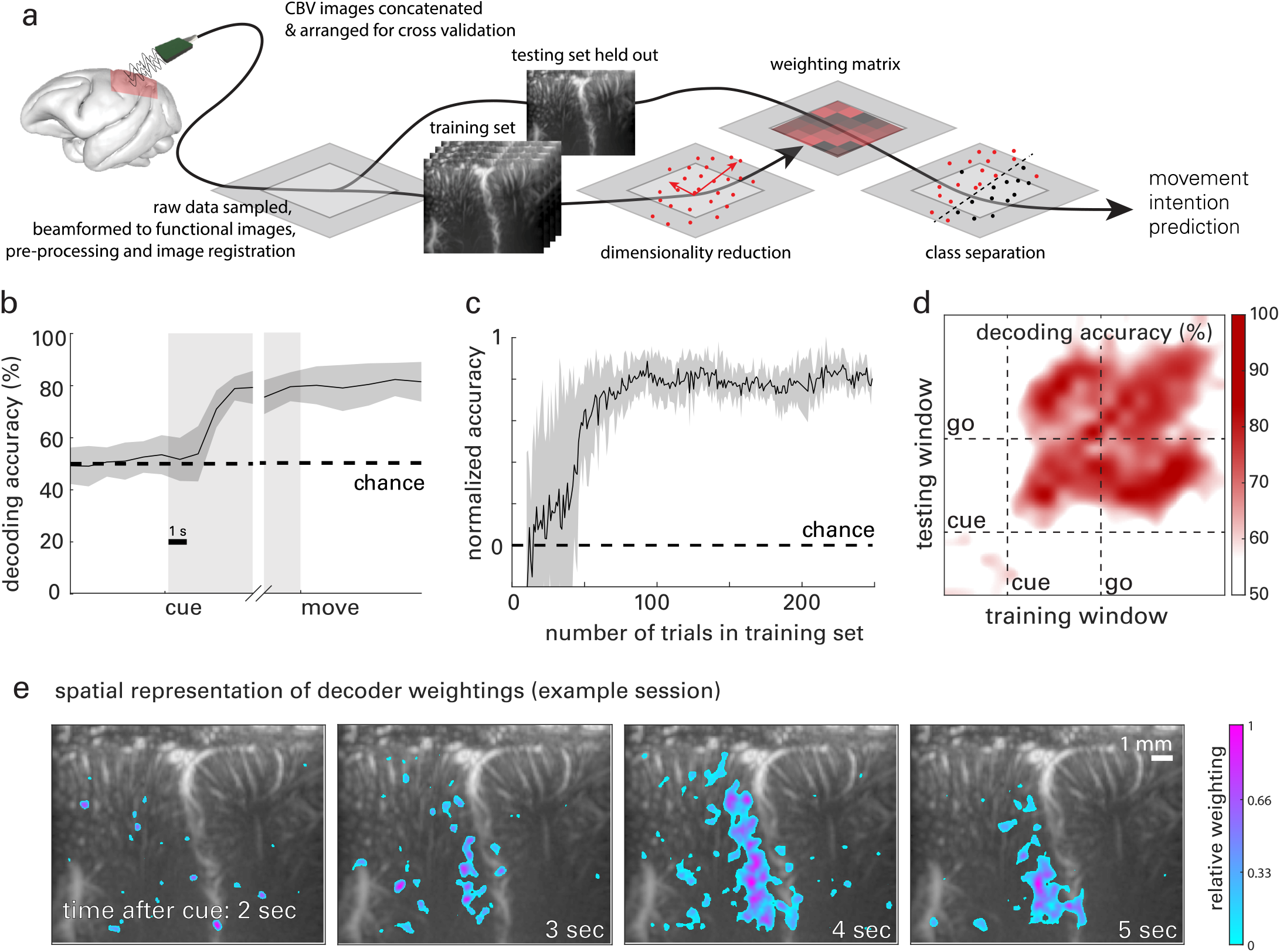
**a**, Data flow chart for cross-validated single-trial movement direction decoding. Training images were separated from testing data according to the cross-validation technique being used. Movement intention predictions were made on a single trial basis based on the dimensionality reduction and classification model built by the training data with corresponding class labels, e.g. actual movement direction. **b**, Decoding accuracy as a function of time across all datasets. **c**, Decoding accuracy as a function of the number of trials used to train the decoder. Data points in b&c are means and shaded areas represent standard error (s.e.m). **d**, Dynamic decoder, i.e. using all combinations of training and testing data, using a one second sliding window. Training the classifiers during the memory or go phase enabled successful decoding of the memory and go phases, suggesting temporally consistent goal information is present in neurovascular dynamics of PPC. **e**, Representative decoder weighting maps. The top 10% most heavily weighted voxels are presented as a function of space and time from 2 to 5 seconds after the go cue was given, overlaid on vascular map.

To analyze the temporal evolution of goal-specific information in PPC, we attempted to decode the movement direction across time through the trial phases: fixation, memory, and movement. For each time point, we used all of the preceding data in both the training and test data. For example, at t = 2 s, we would include all imaging data from t = 0-2 s in the training and test sets. The resulting cross-validated accuracy curves (**Fig. 3b**) show accuracy at chance level during the fixation phase, increasing discriminability during the memory/movement-planning phase, and sustained decode accuracy during the movement phase. That is, the decoder remained at chance level before the monkey received a directional cue, as expected. During the memory phase, decoder accuracy improved, reaching significance 2.08 s +/- 0.82 s after the monkey received the directional cue (but had not yet moved). This accuracy was sustained or increased through the remainder of the memory phase and movement phase.

In a secondary analysis, we removed trials from the training set to determine the amount of data required to achieve maximum decode accuracy (**Fig. 3c**). Within 27 trials, decoder accuracy reached significance for all datasets and continued to increase. In the best case, we could correctly classify movement direction with just 10 trials-worth of training data. Decoder accuracy continued to improve above chance and reached a steady state maximum within 75 trials of training data, on average. This corresponds to roughly 3 minutes of data collection required to achieve meaningful decoding levels and 20 minutes of data collection to achieve nominally maximum levels.

In addition to the encouraging accuracy results, we performed an analysis to determine what information was contained in the brain signals used by the decoder. Were we decoding positions, trajectories, or goals? To answer this question, we used a dynamic decoding technique that uses just 1 s of data to train the decoder. We then attempt to decode the intended direction from 1 s sliding windows of data throughout the trial. Training data and testing data are taken from separate trials using 10-fold cross validation. Finally, we repeat this process, updating the 1 s window used to train the decoder. We do this for all time points through the trial duration. This results in an n x n array of accuracy values where n is the number of time windows tested. The accuracy for each train/decode combination is shown in **Fig. 3d**.

We found high decoding rates across the memory and go phases. This is especially notable in the cases where we decoded movement phase data using decoders trained on memory-delay data and vice-versa. That is, the information encoded by this brain region was highly similar between the memory-delay and go phases. This is evidence that this area is encoding goal information because the movement goal consistent throughout the memory and go phases. This is in contrast to to position or trajectory, which varied across the memory and movement phases within the trial. Thus, information from these parameters would not decode between phases as we saw here. This finding agrees with the canonical function of PPC areas as measured by electrophysiology^31,34^.

Distinct spatial locales within PPC encode goal information, a fact reflected in the variable weighting assigned to each voxel in a decoding algorithm. Roughly speaking, signals with high goal-related SNR are assigned weights of larger magnitude than those with lower SNR and consequently have a larger impact on decoder output. By training decoders that contain a mixed population of PPC neuronal populations, we measured and compared the relative value ascribed to regions, e.g. LIP and PRR, from the perspective of the decoding algorithm. We found that the spatial distributions of the decoder subspaces, i.e. the relative decoder weights across voxels, largely reflected the patterns we saw in the mean activation patterns and statistical maps (example map shown in **Fig. 3e**). That is, the decoder placed highest weightings in area LIP to maximally discriminate the monkey’s goal during the memory period. These weighting likely reflect goal-tuned physiological changes in blood volume induced by neural activity.

### Vascular signal and information content

A purported benefit of the increased spatiotemporal resolution and sensitivity afforded by fUS imaging is access to information content and neurovascular dynamics at scales previously difficult to access with established techniques, such as fMRI. To test these potential benefits, we classified movement goals from the memory phase after synthetically decreasing the resolution of the image. We resized the images using a low-pass filter in each of the dimensions of the imaging plane: x – across the probe surface and z – with image depth. Decoding accuracy continuously decreased as voxel sizes increased (**Fig. 4a**). This effect was isotropic, i.e. similar for both x and z directions. These results underscore the critical importance of fUS’s superior spatial resolution in enabling sophisticated decoding of neural activity.

**Figure 4:**
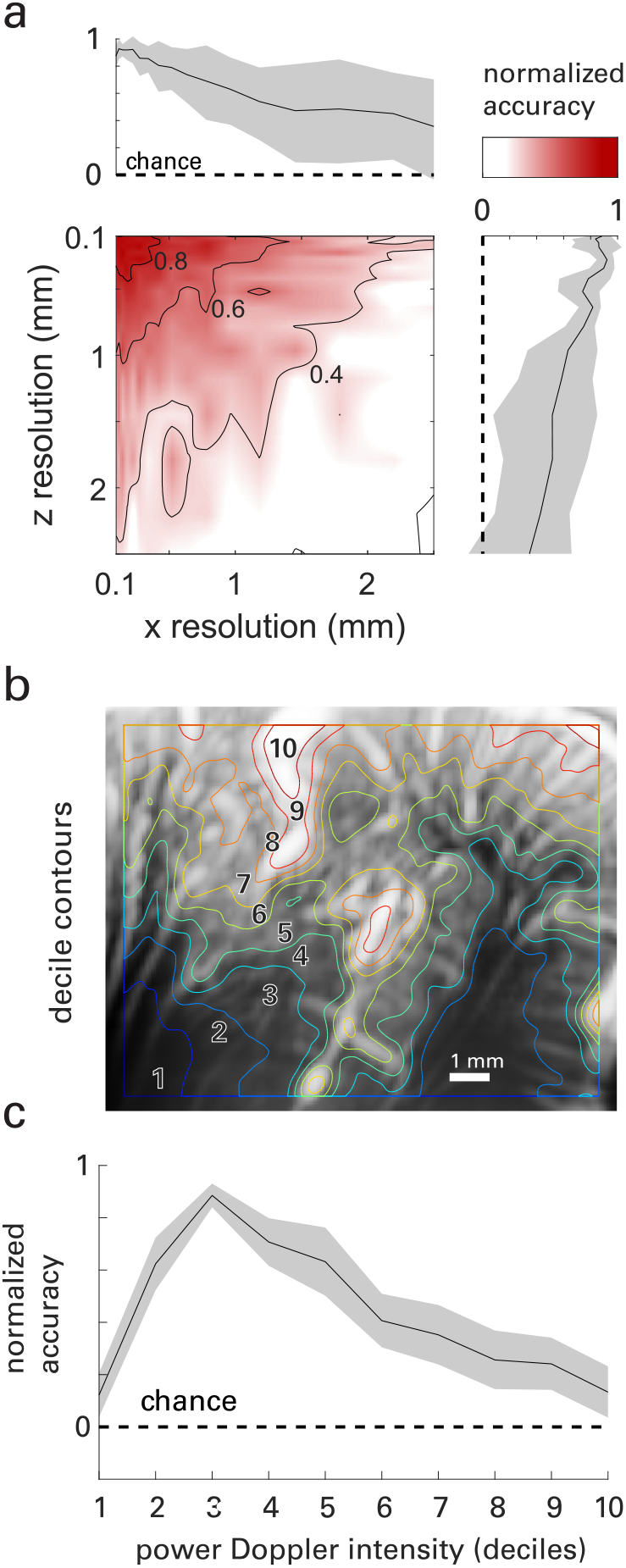
**a**, Decoding accuracy as a function of spatial resolution. Accuracy decreases with resolution in both the x-direction (across the imaging plane) and z-direction (depth in the plane) in an isotropic manner. **b**, A typical vascular map overlaid with contours dividing the image into deciles of mean power Doppler intensity. **c**, Decoding accuracy as a function of the mean power Doppler intensity. Information content is greatest in relatively low-signal areas that correspond to vessel classes that are not individually resolved by fUS. Sub-cortical and primary unit vasculature, i.e. deciles 1 and 10, are least informative to decoding movement direction.

Recent studies show that functional hyperemia starts in a vascular compartment referred to as the primary unit that comprises parenchymal arterioles and first order capillaries, i.e. vessels of diameter < 50 μm^50^. From there, blood flow velocity increases in downstream capillaries and upstream arterioles. We therefore hypothesized that most of the functional information used for decoding would be found in sub-resolution (<100 μm) vessels within the imaging plane. To test this hypothesis, we (spatially) segmented fUS vascular maps of the brain by rank ordering voxels by their mean power Doppler intensity and segmenting them by deciles. A representative spatial map of these ranked deciles is shown in **Fig. 4b**. Whereas deciles 1-2 mostly captured sub-cortical areas, deciles 3-8 mostly captured cortical layers where shallower layers exhibit higher mean intensity. Deciles 9 and 10 were largely restricted to primary unit vasculature, e.g. large arteries, commonly on the cortical surface and in the sulci. We then classified movement goals from individual trials from the memory phase using the rank-ordered voxels, one decile at a time. The resulting decode accuracy as a function of the power Doppler intensity is given in **Fig. 4c**. Accuracy is reported as a fraction of the maximum accuracy for each session. Beginning with the bottom 10% of voxels ordered by mean CBV magnitude, decode accuracies were near chance level. Accuracy increased with increasing Doppler power. Accuracy peaked when the regions of the image within the 3^rd^ decile of mean Doppler power were used to decode movement intention. Finally, accuracy dropped toward chance level as Doppler power increased toward its maximum value for each recording. Thus, goal direction-tuned neurovascular activity, as measured by fUS, exists mostly in vascular anatomy occurring in the cortex. Much of the cortical vasculature is at or below the limits of fUS resolution. Indeed, the 3^rd^ decile regions of cerebral vasculature mostly consists of sub-resolution cortical vessel endings. This result is consistent with the hypothesis that functional hyperemia arises from sub-resolution vessels of the primary unit^20,51^. This finding agrees with previous studies in rodents^52^ and ferrets^24^. These studies showed that the most important contribution to cortical fUS signals were derived from axial flow velocities ranging between 2-10 mm/s, corresponding to vessel diameters <50 μm.

### Memory delayed reaches

To demonstrate the generalizability of fUS BMI, we collected fUS signals while one NHP (monkey H) performed memory reaches via joystick for 6 runs over 4 days. The task was largely similar to that of saccades, but the animal’s gaze remained fixated throughout the trial, including during the fixation, memory, and reach execution phases (**Fig. 5a**). ERAs on the lateral bank of IPS in putative LIP areas reveal populations with direction-specific tuning (**Fig. 5, b-c**). Populations on the medial bank in putative parietal reach region (PRR) do not exhibit such tuning but do show bilateral tuning to the movement (**Fig. 5d**). These results are consistent with electrophysiological recordings, in which the PRR neurons as a population encode both hemispaces, whereas LIP neurons encode largely encode the contralateral space^53^. Decoding accuracy and its temporal evolution was like those observed in the saccade experiments (**Fig. 5e**). Specifically, cross-validated reach goal decodes accuracies ranged from 72.96% (p<0.001) to 94.64% (p<0.001) on a given 30-minute run. The mean accuracy across all sessions and runs was 88.54% (p<0.001).

**Figure 5:**
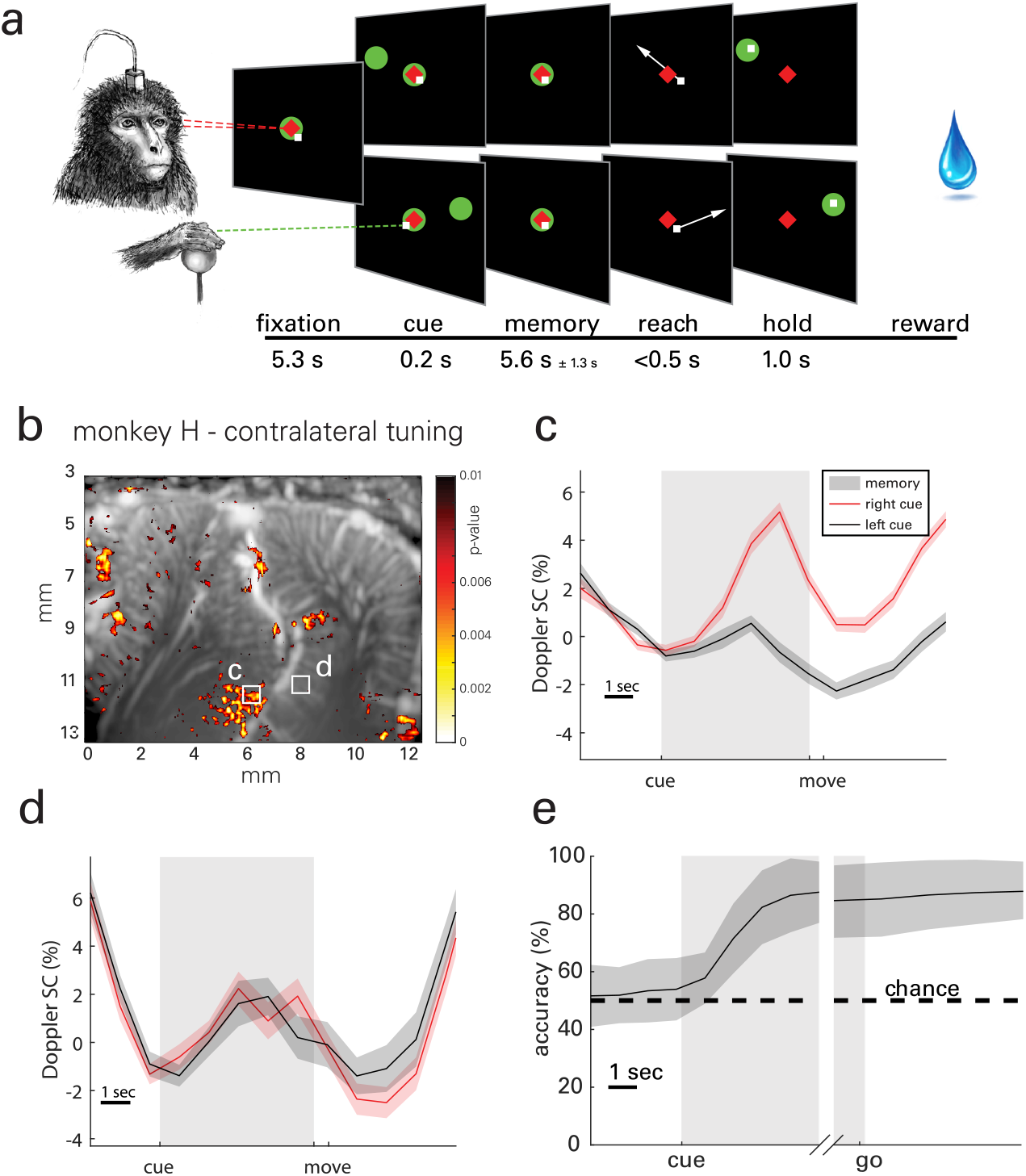
**a**, Memory-guided reaching task. A 2D joystick was positioned in front of the sitting animal, with the handle at the level of his knee. A trial started with the animal fixating on a central cue (red diamond) and positioning the joystick to its center (green circle). Next, a reaching target (green circle) was presented either on the left or the right visual field. The target disappeared after 0.2 s and the animals had to memorize its location while continuing to fixate eye and hand on the center cue. When the hand fixation cue was extinguished (go-signal), the animals performed a reach to the remembered peripheral target location within 0.5 s and maintained the position for another 1 s before receiving a reward. Importantly, eye fixation was maintained through the reaching trial. **b**, A vascular map with ROIs on the lateral and medial banks of the IPS whose ERA waveforms are shown in c and d, respectively. **c**, Waveforms in LIP reveals direction-specific tuning in reaching movements. **d**, Waveforms in the medial bank of IPS exhibit a population with bilateral tuning to reaching movements, consistent with neurophysiological studies in NHPs. **e**, Decoding accuracy as a function of time across all reaching datasets.

## Discussion

Although fUS imaging has been successfully used to monitor brain activity in small (e.g., rodents) and large (e.g., NHP) mammals, the preponderance of previous studies focused on exploring the spatial and temporal responses of behavior across many trials. The present study provides the first direct evidence that fUS imaging is capable of detecting difference in neurovascular response to differing motor goals on a single-trial basis. Two NHPs were trained to perform memory-guided saccade movements to peripheral targets while we recorded fUS activity from the PPC region. One of the animals was also trained to reach to the targets using a joystick. Consistent with previous neurophysiological and brain imaging studies^30–32,35,39^, we found brain areas within PPC that exhibited significant contralateral memory-delayed activity for saccades and reaches. Overall, our findings indicate that the fUS signal from PPC contains information related to planning and execution of motor actions. Importantly, we successfully decoded the intended direction of eye and hand movements on a single-trial basis with significant cross-validated accuracy. This is an essential demonstration for potential application of fUS in developing BMI systems. These results present for the first time a proof of concept that fUS could serve as the basis for brain prosthetics applications, whose portability and minimally invasive nature could vastly expand the user base and applications possible with BMIs.

Electrophysiological studies have also shown that PPC subregions, such as LIP and PRR, can be used as neural basis for the control of oculomotor and reach BMI systems, respectively^46,54–56^. However, there are several very clear advantages of fUS monitoring. The first advantage is a much lower level of invasiveness compared to intracortical electrodes. fUS BMI does not require penetration of the dura mater. This is powerful because penetration of this protective tissue greatly increases the risk level of the implant. Potential infections of the CSF can lead to meningitis, cerebritis, and empyema. In contrast, neurosurgery in the epidural space is significantly less complex and risky, which may better facilitate adoption of epidural approaches. In addition, our approach is highly adaptive thanks to the reconfigurable electronic focusing of the ultrasound beam. This makes it much easier to record from target regions of interest. Electrode arrays are typically inserted only once and can suffer from suboptimal positioning due to poor localization or to avoid piercing major vasculature. Implanted electrode arrays are difficult to reposition as it requires additional surgery. fUS also provides access to cortical areas deep within sulci and subcortical brain structures that are exceedingly difficult to target with electrophysiology. While tissue reactions degrade the performance of chronic electrodes, fUS BMI can, in principle, operate through the dura indefinitely. Fundamentally, the risks of infection, missing targets, and gradual device failure greatly limit the range of scenarios in which intracortical electrical BMIs are considered an option. An epidural fUS BMI would vastly expand the applicable patient populations and research scenarios that could benefit from BMI technology.

A potential limitation of ultrasonic BMI is the temporal latency of the hemodynamic response to neural activity. Increased BMI latency decreases throughput. However, information transfer rate (ITR) can be improved with higher dimensionality decoding and decreased error rates^12,57^. A benefit of fUS imaging is the ability to record the activity of multiple brain areas in parallel. fUS can discriminate responsiveness of neighboring voxels with a functional resolution as fine as 100 μm during auditory stimuli in awake ferrets^24^. The combination of wide field of view and high discriminative resolution provides a very large number of independent voxels for BMI decoders. In the present work, we showed that saccade and reach movements can be decoded from subregions of PPC. Future BMIs can leverage multiple brain areas representing multiple effectors and task variables to increase the dimensionality of the decoder and thus BMI throughput.

The impact of hemodynamic latency also depends greatly on the BMI application being developed. Here, we decoded goals from high-level cognitive areas. Goal decoding circumvents the need for instantaneous updates; goals are formed ahead of an action and remain constant throughout its execution (**Fig. 3d**). While electrophysiology-based BMIs that decode velocity of intended limb movements require as little latency as possible, our results suggest that goal-based decoding is achievable with a fUS based BMI. The development and dissemination of a fUS BMI would be transformative for cognitive BMIs more generally, especially those that operate over large areas of the brain (coverage) on time scales compatible with hemodynamics, including keyboard interfaces or state decoding of mood and other psychiatric states. The extension from 2D fUS to 3D volumetric fUS imaging based on matrix arrays^58^ or row column arrays^59^ would likely improve decoding accuracy^24^ and further expand the impact of fUS for BMI. Finally, there is future potential for faster fUS decoding. For example, a recent study found information present at 20 ms resolution by exploiting fast lag-correlation between neighboring voxels of fUS imaging data in awake and behaving NHPs^26^.

In addition to the technological innovations presented to attain single-trial decoding of movement goals, the data therein were also physiologically meaningful. Typical activation maps and decoder subspaces revealed many active regions within PPC, in scale from ~100 μm to mm to nearly 1 cm, during saccade planning. These responses were more active to contralateral targets; ipsilateral targets did not elicit such activity. The contralateral tuning of these regions during memory-delay is consistent with the findings from previous fMRI studies in our lab, showing that that LIP exhibits stronger changes in BOLD signal for contralateral targets in a similar memory-guided saccade task^39,44^. These results are also consistent with the extensive evidence from electrophysiological recordings in NHPs that show LIP is involved in planning and executing saccadic movements^35,53,60^. Notably, the ERA waveforms display significantly larger target-specific differences compared to fMRI memory-delayed saccade task, i.e., 2-5% vs 0.1-0.5%^39,44^, and have much finer spatial resolution. This sensitivity was possible despite the deepest sub-regions of interest extending >16 mm below the probe surface. At this depth, signal attenuation is approximately −7.2 dB (assuming uniform tissue using a 15 MHz probe). Still, we found significant visuomotor event-related activity (**Fig. 2b**). Using a probe with a lower center frequency would allow for increasing depths at the cost of spatiotemporal resolution. Use of microbubbles^61^ or biomolecular contrast agents^51^ may enhance hemodynamic contrast allowing for deeper imaging without sacrificing resolution.

In addition to well-studied PPC subregions, we also identified patches of activity in the medial parietal area (MP) located within PGm. These functional areas were much smaller in size and magnitude than those nearer to the IPS. This is one potential reason why MP activity was not reported in previous fMRI studies. However, electrophysiological studies have showed that stimulation of this area can elicit goal-directed saccades, indicating its role for eye movements^47^. The addition of the hemodynamic results presented here not only contribute to these findings but are the first hemodynamic evidence of MP function. A limitation of this finding is that we did not observe such activity in the second animal. This is likely due to a targeting problem: we optimized probe positioning to maximally intersect area LIP. Current efforts to develop 3D fUS imaging^58^ will eliminate this limitation, allowing us to identify new areas based on response properties.

We also collected data during a memory-delayed reaching task in one animal. ERA waveforms identified increases in CBV during the memory phase for regions on the medial aspect of the IPS. The parietal reach region (PRR) is located on the medial bank of the IPS and is characterized by functional selectively to effector, i.e. the arm^45^. The responses we observed in this area were indeed effector specific – they did not appear in the saccadic data. However, they were not direction specific, i.e. increases in CBV activity were present for left-cued *and* right-cued trials. This bilateral reach response can be explained by the spatial scale of the recording method. Whereas single unit electrophysiology in PRR reveals single neurons that are tuned to contralateral hand movement planning, a significant portion of PRR neurons are also tuned to ipsilateral movement planning^53^. Within the limits of fUS resolution (~100 μm), each voxel records the neurovascular response to the summed activity of all cells within the voxel (~100 μm x 100 μm x 400 μm). Therefore, our results in the context of previous literature, provide evidence that 1) populations of ipsilaterally- and contralaterally-tuned neurons were roughly equivalent, and 2) these populations are mixed at sub-resolution scales. We also found activity on the lateral bank of IPS that encoded target direction for an upcoming reach. That is, responses were more robust to contralateral targets. Although this area is predominantly involved in saccade movements, neurophysiological studies have also reported neurons within the LIP area that encode reaches to peripheral targets^35,62^.

Approximately 5.6 million US citizens live with paralysis. Paralysis can result from a multitude of conditions including spinal cord injury, stroke, traumatic brain injury, and neurodegenerative disorders. Brain-machine interfacing is a promising technique to restore movement to these populations. The contributions presented here required significant advancements in large-scale recording of neurovascular activity with single-trial sensitivity. These advancements will help to usher in a new era of BMIs that are less-invasive, high-resolution, and scalable. These BMIs will facilitate a direct link between mind and neuroprosthetic devices without damaging healthy brain tissue, significantly minimizing surgical risk. These tools will empower researchers to make unique insights into the function and malfunction of brain circuits. Such insights can be used to improve diagnostic and therapeutic approaches to neurological and psychiatric injury and disease. Furthermore, as future work translates these findings to high-performance minimally invasive BMIs, it will mark a significant advancement in the field of BMIs for people with paralysis.

## Methods

### Animal preparation, implant and probe for fUS imaging in awake behaving monkeys

All surgical and animal care procedures were done in accordance with the National Institutes of Health Guide for the Care and Use of Laboratory Animals and were approved by the California Institute of Technology Institutional Animal Care and Use Committee. We implanted two adult male rhesus macaques (*Macaca mulatta*) weighing 10–13 kg with polyether ether ketone head caps anchored to the skull with titanium screws. We then placed a custom stainless-steel head holder on the midline anterior aspect of the cap. Finally, we placed a unilateral square chamber (2.4 cm inner diameter, Ultem or Nylon) over a craniotomy over the left intraparietal sulcus. The dura underneath the craniotomy was left intact. To guide the placement of the chamber, we acquired high-resolution (700 μm) anatomical MRI images before the surgery using a Siemens 3T MR scanner, with fiducial markers to register the animals’ brains to stereotaxic coordinates. During each recording session, we placed the ultrasound probe (128 elements linear array probe, 15 MHz center frequency, 0.1 mm pitch, Vermon, France) in the chamber with acoustic coupling gel. This enabled us to acquire images from the posterior parietal cortex (PPC) with an aperture of 12.8 mm and depths up to 23 mm. This large field of view allowed us to image several PPC regions simultaneously. These superficial and deep cortical regions included, but were not limited to, area 5d, lateral intraparietal (LIP) area, medial intraparietal (MIP) area, medial parietal area (MP), and ventral intraparietal (VIP) area.

### Behavioral tasks and processing

During each recording session, the monkeys were placed in a dark anechoic room. They sat in a custom designed primate chair, head fixed, facing an LCD monitor ~30 cm away. The animals performed memory-guided eye movements to peripheral targets (**Fig. 2a**). Each trial started with a fixation cue (red diamond; 1.5 cm side length) presented in the center of screen (fixation period). After the animal had fixated for 6.3 s, a single cue (red diamond; 1.5 cm side length) appeared either on the left or the right hemifield for 200 ms, indicating the location of the target. Both targets were located equidistantly from the central fixation cue (23° eccentricity). After the cue offset, the animals were required to remember the location of the targets for a mean of 5.1 s (memory period), while maintaining eye fixation. The memory period varied across sessions from 3.7 to 6.5 s depending on the animals’ training and success rate. Once the central fixation cue disappeared (i.e., go signal), the animals performed a direct eye movement (saccade) within 500 ms to the remembered location of the target. If the eye position arrived within a radius of 5° of the targets, it was reilluminated and stayed on for the duration of the hold period (1 s). If the animal broke eye fixation before the go signal (i.e., shifted their gaze outsize of a window of 7.5 cm, corresponding to 14° of visual angle), the trial was aborted. Successful trials were followed by a liquid reward. The fixation and memory periods were subject to 400 ms of jitter sampled from a uniform distribution to preclude the animal from anticipating the change(s) of the trial phase.

One of the animals (monkey H) also performed memory-guided reach movements to peripheral targets using a 2-dimensional joystick positioned in front of the chair with the handle at knee level. Each trial started with two fixation cues presented at the center of the screen. The animal fixated his eyes on the red diamond cue (1.5 cm side length) and acquired the green cue by moving a square cursor (0.3 cm side length) controlled by the joystick (fixation period). After 5.3 s, a single green target (1.5 cm side length) was presented either on the left or the right visual field for a short period of time (300 ms). After the cue offset, the animal was required to remember the location of the targets for a mean of 5.6 s (memory period), while maintaining eye and hand fixation. The memory period varied across sessions from 4.3 to 6.9 s. Once the central green cue disappeared, the animal performed a direct reach to the remembered target location within 500 ms, without breaking eye fixation. If he moved the cursor to the correct goal location, the target was re-illuminated and stayed on for duration of the hold period (1 s). Targets were placed at the same locations as in saccade trials. If the cursor moved out of the target location, the target was extinguished, and the trial was aborted. Any trial in which the animal broke eye fixation or initiated a reaching movement before the go signal or failed to arrive at the target location was aborted. Successful trials were followed with the same liquid reward as in saccade trials.

Visual stimuli were presented on an LCD monitor using custom Python software based on PsychoPy^63^. Eye position was monitored at 60 Hz using a miniature infrared camera (Resonance Technology, Northridge, CA, USA) and ViewPoint pupil-tracking software (Arrington Research, Scottsdale, AZ, USA). Reaches were performed using a 2-dimensional joystick (Measurement Systems). Both eye and cursor positions were recorded simultaneously with the stimulus and timing information and stored for offline access. Data analysis was performed in Matlab 2019b (MathWorks, Natick, MA, USA) using standard desktop computers.

### Functional Ultrasound (fUS) sequence and recording

We performed all acquisitions with a single 128-element piezoelectric ultrasound probe with a center frequency of 15.6 MHz. Scanning was performed with an ultrafast ultrasound research scanner (Vantage 128 by Verasonics, Kirkland, WA). To image functional changes in the brain, we used ultrafast ultrasound Doppler sequences. As the source of contrast in Doppler images originates from red blood cell motion, regional changes in power Doppler intensity are proportional to cerebral blood volume (CBV) changes induced by neurovascular coupling^17^. Specifically, our sequence transmitted plane waves at five discrete angles: −6, −3, 0, 3, and 6 degrees. We then coherently compounded the radiofrequency (RF) data acquired along each angle into a single image with higher contrast. We acquired sets of 250 coherently compounded frames at 500 Hz every second (500 ms period of acquisition and 500 ms of rest).

Anatomical PPC regions were spatially located by their stereotaxic positions from the pre-surgical MRI. Response of these functional areas was confirmed by mapping activated voxels obtained during the experimental phase of this work. If necessary, the imaging plane was adjusted to record the most responsive area. Each acquisition consisted of 900-3600 blocks of 250 frames where each block represented 1 second of data (equivalent to 15-60 minutes runtime). Finally, we stored the in-phase and quadrature sampled data to high-speed solid-state drive memory for offline processing.

### Beamforming and Doppler signal processing

We used singular value decomposition to discriminate red blood cell motion from tissue motion and extracted the Doppler signal in each ensemble of 250 coherently compounded frames^64,65^. The resulting images were then stored in a 3D array of 2D images in time series. In some experiments, we observed motion of the entire imaging frame. These shifts were indicative of a change in the position of the probe/tissue interface due to uncommonly forceful movements of the animal. We corrected for these events using rigid-body image registration based on the open source NoRMCorre package^66^, using an empirical template created from the first 20 frames from the same session. We also tested nonrigid image registration but found little improvement, confirming that motion observed was due to small movements between the probe/dura interface rather than changes in temperature or brain morphology.

We display event-related average (ERA) waveforms (**Fig. 2, c-f,h-i; Fig. 5, c-d**) of power Doppler change as percentage change from baseline. The baseline consists of the three seconds preceding the first Doppler image obtained after the directional cue was given on any given trial. ERA waveforms are represented as a solid line with surrounding shaded areas representing the mean and standard deviation. We generated activation maps (**Fig. 2, a,g**) by performing a two-sample t-test for each voxel individually with false discovery rate (FDR) correction based on the number of voxels tested. In this test, we compared the area under the curve of the change in power Doppler during the memory phase of the event-related response. The movement direction represented the two conditions to be compared, and each trial represented one sample for each condition. Voxels with values of p<0.01 are displayed as a heat map overlaid on a background vascular map for anatomical reference.

### Single trial decoding

Decoding single trial movement intention involved three parts: 1) aligning CBV image time series with behavioral labels, 2) feature selection, dimensionality reduction and class discrimination, and 3) cross validation and performance evaluation (**Fig. 3a**). First, we divided the imaging dataset into event aligned responses for each trial, resulting in a 4-dimensional data-task structure, i.e. 2D CBV images through time for each trial. We then separated trials into a training set and testing set according to a 10-fold cross validation scheme. The training set was attached to class labels that represented movement direction (i.e. Left vs. Right); the test set was stripped of such labels. Features were selected by ranking each voxel’s t-value comparing the memory-phase responses to target direction. Direction-tuned voxels (FDR corrected, q<0.05) up to 10% of the total image were kept as features. For dimensionality reduction and class separation, we used classwise principal component analysis (CPCA) and linear discriminant analysis (LDA)^48^, respectively. This decoding method has been implemented with success in many real-time BMIs^16,67–70^, and is especially useful in applications with high dimensionality. CPCA computes the principal components (PCs) in a piecewise manner individually for training data of each class. We retained principal components to account for >95% of variance. We improved class separability by running linear discriminant analysis (LDA) on the CPCA-transformed data. Mathematically the transformed feature for each trial can be represented by *f* = *T_LDA_*Φ_*CPCA*_(*d*), where *d* ∈ ℝ^1^ are the flattened imaging data for a single trial, Φ_*CPCA*_ is the piecewise linear CPCA transformation, and *T_LDA_* is the LDA transformation. Φ_*CPCA*_ is physically related to space and time and thus can be viewed within the context of physiological meaning (**Fig. 3e**). We subsequently used Bayes rule to calculate the posterior probabilities of each class given the observed feature space. Because CPCA is a piecewise function, this is done twice, i.e. once for each class, resulting in four posterior likelihoods, i.e. *P_L_*(*L|f*\*), *P_L_*(*R|f*\*), *P_R_*(*L|f*\*), *P*(*(R|f*\*), where *f*\* represents the observation, *P_L_* and *P_R_* represent the posterior probabilities in the CPCA subspaces created with training data from left-directed and right-directed trials, respectively. Finally, we store the optimal PC vectors and corresponding discriminant hyperplane from the subspace with the highest posterior probability. We then used these findings to predict movement direction for each trial in the testing set. That is, we compute *f*\* from fUS imaging data for each trial in the testing set to predict the upcoming movement direction. Finally, we rotate the training and testing sets according to k-fold validation, storing the BMI performance metrics for each iteration. We report the mean decoding accuracy as a percentage of correctly predicted trials (**Fig. 3b**). In measures across multiple sessions where an independent variable is being tested (e.g. number of trials in training set), we use a normalized accuracy that is linearly scaled to [0, 1] where 0 is chance level, i.e. 50%, and 1 is the maximum accuracy across the set of values used in the independent variable (e.g. **Fig. 3c**). This was necessary to account for significant differences in raw accuracy values across multiple sessions and animals.

As BMI models increase in complexity, their need for data also increases. To demonstrate the robustness of our piecewise linear decoding scheme to limited data, we systematically reduced the amount of data used in the training set (**Fig. 3c**). We used *N-i* trials in the training set and *i* trials in the testing set in a cross-validated manner, rotating the training/testing set *i* times for *i* = 1, 2,… *N-10*. We stop at *N-10* because accuracy was diminished to chance level and when less than 10 trials are used in the training set, it becomes increasingly likely that there will be an under- or non-represented class, i.e. few or no trials to one of the movement directions. We report the mean normalized accuracy standard error of the means (SEM) across both animals and all recording sessions as a function of the number of trials in the training set (*N-i*) (**Fig. 3c**).

We also performed an analysis to determine the nature of hemodynamic encoding in PPC using a dynamic decoding technique. In this analysis, we used all temporal combinations of training and testing data, using a one second sliding window. We used 1 s of data from all trials to train the decoder and then attempted to decode from each of the 1 s windows of testing data throughout the trial. We then updated the training window and repeated the process. This analysis results in an *n x n* array of accuracy values where *n* is the number of time windows in the trial. We report the 10fold cross-validated accuracies as the percentage of correctly predicted trials (**Fig. 3d**).

Part of the motivation for using fUS is its spatial resolution. To test the effects of increased resolution, we synthetically reduced the resolution of the in-plane imaging data using a Gaussian filter. We performed this analysis at all combinations of x and z direction (width and depth, respectively) starting at true resolution, i.e. 100 μm, up to a worst-case of 5 mm resolution. We report the 10-fold cross-validated accuracy values as a function of these decreasing resolutions as a 2D heat map and as 1D curves of mean accuracy in both the x and z directions with shaded areas representing SEM (**Fig. 4a**). A limitation of this approach is that we cannot downsample the out-of-plane dimension. Thus, the reported accuracy values are likely higher than those attainable by a technique with isotropic voxel size, e.g. fMRI.

We also investigated the source of neurovascular information content by segmenting the images according to their mean power Doppler signal as a proxy for mean cerebral blood flow within a given area. Specifically, we segmented the image into deciles by mean power Doppler signal within a session, where higher deciles represented higher power and thus higher mean blood flow (**Fig. 4b**). Deciles were delineated by the number of voxels, i.e. the number of voxels was the same within each segment and did not overlap. Using only the voxels within each decile segment, we computed the mean accuracy for each recording session. We report the mean normalized accuracy across all recording sessions (**Fig. 4c**) where shaded error bars represent SEM.

## Acknowledgements

We thank Kelsie Pejsa for assistance with animal care, surgeries, and training. We thank Thomas Deffieux for his contributions to the ultrasound neuroimaging methods that made this work possible. We also thank Igor Kagan for assisting in MR scanning and chamber placement planning. Finally, we thank Krissta Passanante for her illustrations. DM was supported by a Human Frontiers Science Program Cross-Disciplinary Postdoctoral Fellowship (Award No. <u>LT000637/2016</u>). This research was supported by the National Institute of Health BRAIN Initiative (grant <u>U01NS099724</u> to MGS), the T&C Chen Brain-machine Interface Center, and the Boswell Foundation.

## Contributions

S.L.N., D.M., V.N.C., M.T., M.G.S. and R.A.A. conceived the study; S.N., D.M., C.D., and M.T. developed the imaging sequences; S.L.N., V.N.C., and W.G. trained the animals. S.L.N., V.N.C., D.M. and W.G. acquired the data; S.L.N., D.M. and V.N.C. performed the data processing; S.L.N., D.M., and V.N.C. drafted the manuscript with substantial contribution from M.G.S. and R.A.A.; all authors edited and approved the final version of the manuscript. M.T., M.G.S. and R.A.A. supervised the research.

